# The Gb3-enriched CD59/flotillin plasma membrane domain regulates host cell invasion by *Pseudomonas aeruginosa*

**DOI:** 10.1101/2020.06.26.173336

**Authors:** Annette Brandel, Sahaja Aigal, Simon Lagies, Manuel Schlimpert, Anika Lehmann, Daniel Hummel, Daniel Fisch, Ana Valeria Meléndez, Josef Madl, Thorsten Eierhoff, Bernd Kammerer, Winfried Römer

## Abstract

The opportunistic pathogen *Pseudomonas aeruginosa* is responsible for a high number of acute and chronic hospital-acquired infections. As it develops more and more resistances against existing antibiotics, *P. aeruginosa* has been placed highest on the global priority list of antibiotic-resistant bacteria for which alternative treatments are urgently needed. Former studies have highlighted the crucial role of the bacterial lectin LecA and the host cell glycosphingolipid globotriaosylceramide (Gb3) for the cellular uptake of *P. aeruginosa* into epithelial cells via the lipid zipper mechanism. To further characterize the host cell plasma membrane domain for LecA-driven attachment and invasion, we analyzed the protein and lipid composition of pulled-down membrane domains for novel interaction partners of LecA by mass spectrometry. We unraveled a predilection of LecA for binding to saturated Gb3 species in the extracellular membrane leaflet and an induction of dynamic phosphatidylinositol (3,4,5)-trisphosphate clusters at the intracellular leaflet co-localizing with sites of LecA binding. Moreover, we identified the GPI-anchored protein CD59 and flotillins, known as cargo and eponymous component of flotillin-assisted endocytosis, as LecA interaction partners. Depletion of each of these host cell proteins resulted in more than 50% of reduction in invasiveness of the *P. aeruginosa* strain PAO1 highlighting the importance of this LecA-induced plasma membrane domain. Our strategy to reduce the complexity of host-pathogen interactions by first identifying interaction partners of a single virulence factor and subsequently transferring these findings to the bacterium has been proven to be a successful approach in elucidating the molecular mechanisms of bacterial infections.

## 1. Introduction

*Pseudomonas aeruginosa* (PA), a multi-drug resistant bacterium takes a spot on WHO’s highest priority list [1]. This pathogen infects lungs, skin wounds and burns as well as the urinary and gastrointestinal tract of immuno-compromised individuals [2,3]. The very first contact between a pathogenic bacterium and its host cell can already decide the course of infection. Proteins aiding bacteria to successfully colonize epithelial cells are broadly classified into adhesins and invasins. The two lectins of PA, namely, LecA and LecB, were initially defined as exemplary adhesion proteins [4], however, several reports now suggest an additional role of LecA in the pathogenicity of PA [5–8]. The homotetrameric protein is localized to the outer membrane of the bacterium [9] and preferentially binds to the glycosphingolipid (GSL) globotriaosylceramide (Gb3; also known as CD77 or the P^k^ blood group antigen). Gb3 is well known to be the receptor for the B-subunit of AB_5_ toxins produced by *Shigella dysenteriae* and enterohemorrhagic strains of *Escherichia coli* (Shiga toxin and Shiga-like toxin B-subunit, respectively, here referred to as StxB [10]). We previously demonstrated the importance of the interaction between Gb3 and LecA for PA uptake into lung epithelial cells [8]. Moreover, a novel divalent LecA ligand identified from a galactoside-conjugate array, bound to the lectin with high affinity and lowered the invasiveness of PA by up to 90% [11].

Pathogens hijack GSLs to induce plasma membrane bending and to transduce signals promoting endocytosis [12,13]. Both PAO1 and its lectin LecA trigger the activation of CrkII, an adaptor protein implicated in various cellular processes including cell adhesion and cytoskeletal reorganization [14,15], by phosphorylation of CrkII at Tyr^221^ [16,17]. Interestingly, induction of this signaling pathway was dependent on binding of LecA to Gb3 located in the extracellular leaflet of the plasma membrane. Clearly, these observations raise the question of how the signal generated from receptor binding at the extracellular membrane leaflet, is transmitted to the intracellular site, activating proteins such as CrkII. One possibility could be the involvement of GSLs with long fatty acyl chains interdigitating into the inner leaflet of the membrane bilayer [18–20]. Additionally, the local lipid and protein environment of Gb3 may affect the induced signaling events and endocytic trafficking routes. The lipid environment as well as length and saturation level of the fatty acyl chains of GSLs influence orientation and accessibility of the carbohydrate groups and affect the binding behavior of StxB to Gb3 in model membrane systems [21–23]. Ordered domains in the outer leaflet of the cellular plasma membrane are often termed lipid rafts. These domains are enriched in GSLs, cholesterol and glycosylphosphatidylinositol (GPI)-anchored proteins and serve as sorting and signaling platforms [24–26]. Due to the high saturation level of the fatty acyl chains of Gb3, the lipid is preferentially, but not exclusively located in lipid rafts [23,27,28].

Many bacterial species evade the immune system and invade the target host cell. The pathogen entry mechanism and its virulence factors comprise an astonishing variety of possibilities. The best-characterized endocytic pathways include clathrin- and caveolin-mediated uptake [24,25]. Inhibition of these pathways led to the discovery of clathrin- and caveolin-independent uptake mechanisms including flotillin-assisted endocytosis [31]. Flotillins are heterotetrameric protein complexes consisting of flotillin-1 and flotillin-2 and known to scaffold lipid rafts [32,33]. Flotillins anchor to the cytosolic face of the plasma membrane and to endosomal structures via myristoylation and palmitoylation, and have been implicated in cell adhesion, vesicle trafficking and cytoskeleton rearrangement [34–36]. Importantly, flotillin-enriched domains also represent active cellular signaling platforms typically involving Src family kinases [37,38]. A role of flotillins in endocytosis was first described for the GPI-anchored complement inhibitor protein CD59 and the GSL GM1, the receptor for cholera toxin [31], and later also for the amyloid precursor protein, glutamate- and dopamine-transporters [39,40].

The search for novel therapeutic approaches for multi-drug resistant bacteria is driven by the complexity of bacterial infections, host-pathogen interaction dynamics and the rapid adaptation capacity of bacteria. In this study, we focus on the virulence factor LecA to better understand initial processes of host-pathogen interactions. So far, little is known about important players within and associated to the Gb3-enriched plasma membrane domain leading to the induction of endocytosis and signaling of PA through its lectin LecA. Here, we characterize both, proteins and lipids of this membrane domain and identify interaction partners of LecA and Gb3 using a pull-down strategy followed by mass spectrometry (MS) analysis. We unravel an involvement of CD59, flotillins, Src family kinases and phosphatidylinositol (3,4,5)- trisphosphate (PIP_3_) in the process of LecA binding to the plasma membrane. As major finding, we demonstrate that CD59 and flotillins promote PA invasion into lung epithelial cells. Deepening our understanding of the interplay between virulence factors and the host cell plasma membrane is crucial to develop novel treatment strategies and to minimize the rising risk of untreatable bacterial infections.

## 2. Materials and methods

### Cell culture and lectin stimulation

The human lung epithelial cell line H1299 (American Type Culture Collection, CRL-5803) was cultured in Roswell Park Memorial Institute (RPMI) medium supplemented with 10% fetal calf serum (FCS) and 2 mM L-glutamine at 37 °C and 5% CO_2_. LecA was expressed in *E. coli* BL21 (DE3), transformed with the plasmid pET25-pa1l encoding the lectin and purified as published in [7]. The B-subunit of Shiga toxin 1 (StxB) was bought from Sigma Aldrich. Lyophilized StxB was resuspended in ultrapure water, while LecA was diluted in calcium- and magnesium-free Dulbecco’s phosphate-buffered saline (DPBS−/−). Both lectins were filtered sterile and stored at 4 °C. Cells were seeded in appropriate amounts and stimulated at a confluency of 70-80% with a final concentration of 100 nM LecA or StxB for indicated time points. For cholesterol depletion, cells were pre-treated with 10 mM methyl-beta-cyclodextrin (MβCD) in RPMI for 30 min at 37 °C and washed once with DPBS−/− before lectin stimulation.

### SDS-PAGE and immunoblot analysis

Stimulated H1299 cells were harvested in radio-immunoprecipitation assay (RIPA) buffer (20 mM Tris (pH 8.0), 0.5% (w/vol) sodiumdeoxycholate, 13.7 mM NaCl, 10% (vol/vol) Glycerol, 0.1% (w/vol) SDS, 2 mM EDTA, freshly supplemented with phosphatase and protease inhibitors (Sigma)) for 60 min at 4 °C. Protein concentrations were determined by Pierce BCA Protein Assay Kit (Thermo Fisher Scientific). Cell lysates were denatured in SDS loading buffer and boiled at 95 °C for 7 min. Equal amounts of proteins were separated on 8 or 12% Tris-glycine gels and transferred on a nitrocellulose membrane by application of 0.12 A/gel. BSA (3% in TBS-T (50 mM Tris (pH 7.6), 154 mM NaCl, 0.5% (vol/vol) Tween 20)) was used to inhibit unspecific binding to the membrane before primary antibodies (1:1000) were added at 4 °C overnight. The corresponding horseradish peroxidase-conjugated secondary antibody (1:2000) was incubated with the membrane for another hour at room temperature (RT) the following day. Finally, ClarityTM Western ECL Chemiluminescent Substrate (Bio-Rad) was used to detect the luminescence signal by a Vilber Lourmat Fusion FX chemiluminescence imager. For the detection of LecA-biotin, an IRDye 800CW Streptavidin antibody (926-32230) was used and fluorescence signal was measured by the Odyssey CLx (LI-COR). Protein levels were analyzed by means of Fiji ImageJ 1.0 software and gel analysis tools.

### Pull-down and immunoprecipitation

For pull-down studies, lectins were biotinylated by NHS-ester conjugation (Thermo Fisher Scientific) and dialyzed against DPBS−/− overnight. H1299 cells treated with 100 nM LecA-biotin or StxB-biotin for indicated time points were lysed by a buffer composition consisting of 25 mM Tris/HCl (pH 7.4), 150 mM NaCl, 1 mM EDTA, 1% NP40 and 5% glycerol for 60 min at 4 °C. After a short clarification centrifugation step, normalized protein lysates were incubated with magnetic streptavidin beads (Thermo Fisher Scientific) for 3 h at 4 °C. Subsequently, beads were washed three times with lysis buffer and in between rotated for 10 min at 4 °C. Pulled-down proteins were eluted by 2x SDS loading buffer and subjected to immunoblot analysis for further characterization. For co-immunoprecipitation, the Capturem Protein A technology from Takara was employed according to the manufacturer’s protocol. Briefly, H1299 cells were stimulated with LecA as described above and lysed by the provided lysis buffer freshly supplemented with protease inhibitor cocktail. Cell lysates were incubated with the target primary antibody for 60 min at 4 °C and end-to-end rotation. The antibody/antigen complex was loaded on equilibrated spin columns, washed and eluted in elution buffer. Finally, 5x SDS loading buffer was added to the eluate and samples were boiled and analyzed by immunoblot analysis as described. For depletion of GSLs, cells were cultivated four days in the presence of 2.5 μM D-threo-l-phenyl-2-palmitoylarmino-3-morpholino-l-propanol (PPMP; Santa Cruz) to inhibit synthesis of glucosylceramide-based GSLs [41].

### Lipid analysis by MS

Untreated or MβCD pre-incubated H1299 cells were stimulated with biotinylated lectins for 20 min at 37 °C. Pull-down was performed as described above with the exception that streptavidin beads were finally eluted in methanol/water (1:1) and stored at −80 °C until further usage. Lipid extraction by centrifugation with chloroform was performed as previously described [42]. Lipids were reconstituted in isopropanol:acetonitrile:water 2:1:1. The Gb3 composition and binding affinity of the cells were assessed by high-resolution mass spectrometry (Synapt G2Si, Waters Corporation) operated in positive MS^E^ mode. Cholesterol and its influence on Gb3 binding were analyzed by targeted LC-QqQ-MS (6460, Agilent Technologies) operated in SRM/MRM mode. Chromatographic separation was achieved on a BEH C18 column (100 mm × 2.1 mm, 1.8 μm, Waters Corporation) as previously described [43]. For evaluation of the lectin preferences, pulled-down Gb3 species were normalized to their input values (i.e. the total cell lysates before incubation with streptavidin-coated beads). Additionally, to correct for different binding affinities, numbers were normalized to the species C16:0.

### Protein analysis by MS

The pull-down of H1299 cells stimulated with LecA for 5 and 15 min was conducted as described above. After incubation of the cell lysates with magnetic streptavidin beads, beads were washed three times with lysis buffer. Further processing for protein MS was performed at Toplab GmbH in Planegg-Martinsried. The samples were submerged in cleavage buffer (8 M urea/ 0.4 M NH_4_HCO_3_ buffer) and reduced with 5 μl 45 mM dithiothreitol for 30 min at 55 °C before they were alkylated with 5 μl 100 mM iodoacetamide for 15 min at RT in the dark. For trypsin digestion, the samples were diluted to 2 M urea / 0.1 M NH_4_HCO_3_ with 140 μl of HPLC grade water (VWR). Digest with mass-spec grade trypsin (Serva, porcine) was performed at 37 °C overnight. For nano-LC-ESI-MS/MS the digests were acidified to 0.05% FA and 10 μl of the digests were subsequently injected. HPLC separation was done using an EASY-nLC1000 (Thermo Scientific) system with the following columns and chromatographic settings: The peptides were applied to a C18 column (Acclaim PepMap 100 pre-column, C18, 3 μm, 2 cm × 75 μm Nanoviper, Thermo Scientific) and subsequently separated using an analytical column (EASY-Spray column, 50 cm × 75 μm ID, PepMap C18 2 μm particles, 100 Å pore size, Thermo Scientific) by applying a linear gradient (A: 0.1% formic acid in water, B: 0.1% formic acid in 100% ACN) at a flow rate of 200 nl/min. The gradient used was: 1-25% B in 120 min, 25-50% B in 10 min, 84% B in 10 min.

MS analysis was conducted on a LTQ Orbitrap XL mass-spectrometer (Thermo Scientific), which was coupled to the HPLC-system. The mass spectrometer was operated in the so-called “data-dependent” mode where after each global scan the five most intense peptide signals are chosen automatically for MS/MS-analysis.

The LC-ESI-MS/MS data were used for a database search with the software Mascot (Matrix Science) using the SwissProt database, species: human. Peptide mass tolerance was set to 50 ppm, fragment mass tolerance was set to 0.6 Da, a significance threshold p<0.05 was used. Carbamidomethylation at cystein was set as fixed modification; oxidation at methionine was set as variable modification. Peptides with up to 1 missed cleavage site were searched. Identified proteins are sorted by the Exponentially Modified Protein Abundance Index (emPAI), which offers approximate, label-free, relative quantitation of the proteins in a mixture based on protein coverage by the peptide matches in a database search result [44].

### Lectin labeling and immunofluorescence microscopy

Alexa Fluor-488 or Alexa Fluor-647 dyes (Thermo Fisher Scientific) were used to label the lectins according to the manufacturer’s protocol. Hereby, dyes were used in a fivefold molar excess. H1299 cells were seeded on glass coverslips and allowed to adhere. The next day, cells were stimulated with fluorescently labeled LecA for 60 min. For the inhibition of phosphoinositide 3-kinase (PI3)-kinases, cells were pre-treated with 100 nM Wortmannin (Sigma-Aldrich) for 30 min and stimulated with LecA. Subsequently, cells were fixed with 4% formaldehyde for 15 min at RT and/or subjected to ice-cold methanol for 8 min at −20 °C. The membrane was permeabilized by 0.2% Triton-X 100 in DPBS−/−, if the cells were not treated with methanol. Fixed cells were blocked in 3% BSA in DPBS−/− for 30 min and stained for target proteins by incubation with primary antibodies (1:100) for 1 h at RT. After three washes, cells were stained with fluorescently-labeled secondary antibodies (1:200) for 30 min at RT in the dark. Subsequently, the nucleus was counterstained with DAPI (1:1000) and the samples were mounted on cover slips using Mowiol (containing the anti-bleaching reagent DABCO). Samples were imaged by means of a laser scanning confocal microscope system from Nikon (Eclipse Ti-E, A1R), equipped with a 60x oil immersion objective and a numerical aperture of 1.49. Co-localization was calculated in Fiji ImageJ 1.0 software using the Coloc2 plugin. A minimum of three biological replicates and five to ten areas (one to five cells) per condition and replicate were analyzed.

### G-LISA

Rac1 activation was measured by means of a G-LISA Activation Assay with luminescence read-out (Cytoskeleton, Inc.) according to the manufacturer’s protocol. Briefly, LecA-treated and snap frozen cell lysates were subjected to a 96 Rac-GTP affinity well plate and incubated for 30 min. Following a 2 min incubation with antigen presenting buffer, anti-Rac1 antibody was added and the plate was vigorously shaken for 45 min at RT. The procedure was repeated with a corresponding secondary antibody. Subsequently, the luminescence signal was detected using a HRP detection reagent and a microplate reader (Synergy H4, Biotek).

### Plasmid transfection

H1299 cells were transfected with 1 μg plasmid for single transfections and 0.5 μg for co-transfection of two plasmids. Lipofectamine 2000 was used as transfection reagent (Thermo Fisher Scientific). Cells were kept in Opti-MEM Reduced Serum Media (Thermo Fisher Scientific) during the 3 h incubation before medium was exchanged to RPMI. The plasmid pcDNA3-AKT-PH-GFP (Addgene #18836 (Craig Montell)) was requested from the Signaling Factory of the Albert-Ludwigs-University Freiburg. Flotillin-1- and flotillin-2-mCherry were kind gifts from A. Echard (Institut Pasteur, Paris, France).

### Live-cell imaging

H1299 cells were grown on glass cover slips (Thermo Fisher Scientific) until 80% confluency. On the day of the experiment, cells were pre-washed with and kept in Hanks’ Balanced Salt Solution (HBSS, Thermo Fisher Scientific) while imaging. Live cell imaging was performed at 37 °C by using an incubator stage (OKOLab) mounted onto a confocal laser scanning microscope (Nikon Eclipse Ti-E, A1R).

### Creation of Δ*FLOT1* cell lines with CRISPR/Cas9 system

Knockout of *Flotillin-1* was accomplished according to the protocol previously described [45]. Briefly, single guide RNAs (sgRNAs, designed using crispr.mit.edu, listed in Table S1) were cloned into pX458 vector (Addgene plasmid #48138) and transfected into H1299 cells. After 72 h, the transfected cells were trypsinized and resuspended in FACS buffer. GFP positive cells were sorted into 96-well plates. After two weeks, single cell clones were expanded into bigger culture dishes. The initial screening for knockout clones was conducted by immunoblotting against flotillin-1 antibody. The genomic locus was amplified by PCR and analyzed by Sanger sequencing in order to confirm the knockout.

### Quantitative RT-PCR

The single clones were subjected to TRIzol (Sigma-Aldrich) treatment in order to extract mRNA. Subsequently, the extracted mRNA was transcribed into cDNA using the Maxima First Strand cDNA Synthesis kit. SYBR Select Master Mix for CFX was used for qPCR according to the manufacturer’s protocol. Samples were analyzed using the CFX384 qPCR system (Biorad, version 3.0). Relative flotillin mRNA levels were normalized to GAPDH (glyceraldehyde 3-phosphate dehydrogenase) mRNA levels. Primer sequences are listed in Table S1.

### siRNA transfection

H1299 cells were depleted by means of on-target SMART pools of siRNA against flotillin-2 (Cat. No. L-003666-01-0010) or CD59 (L-004537-02-0005) purchased from Dharmacon, Horizon Discovery. During the incubation with a mixture of siRNA and Lipofectamine 2000, cells were kept in Opti-MEM Reduced Serum Media. Sequences of the siRNA are listed in Table S1.

### Invasion assay

*P. aeruginosa* strain PAO1 was cultivated, GFP-tagged and deleted of LecA as described before [8]. For the invasion assay, overnight cultures of PAO1 wild-type (WT) and ΔLecA were centrifuged before resuspension of the pellet in RPMI containing 1 mM CaCl_2_ and MgCl_2_. H1299 cells (control siRNA (Qiagen), CD59- or flotillin-depleted) were treated with PA at a multiplicity of infection (MOI) of 100 for 2 h at 37 °C. After washing with PBS, cells were treated with amikacin sulphate (400 μg/ml; Sigma-Aldrich) for 2 h at 37 °C to exclude extracellular bacteria. Finally, the cells were lysed with 0.25% (vol/vol) Triton X-100, plated on LB-Agar plates containing 60 μg/ml Gentamicin and incubated at 37 °C overnight. The following day, colonies were counted and invasion rate was calculated as percentage of Amikacin-survived bacteria to the total number of bacteria not treated with Amikacin. Mean values of 3 individual experiments were normalized to the invasion rate of WT PAO1 into WT H1299.

### Antibodies and chemical reagents

The following antibodies were obtained from commercial sources: monoclonal mouse anti-CD59 (MEM-43, Abcam, Cat. No. ab9182), monoclonal rabbit anti-flotillin-1 (D2V7J) XP (Cell Signaling, Cat. No. 18634), monoclonal mouse anti-flotillin-2 (BD Biosciences, Cat. No. 610383), polyclonal rabbit anti-GAPDH (Sigma, Cat. No. G9545), monoclonal rabbit anti-phospho-Src Family (Tyr^416^, Cell Signaling, Cat. No. 6943), monoclonal rabbit anti-Src Family (Cell Signaling, Cat. No. 2109). LecA was detected by a custom-made polyclonal rabbit anti-LecA antibody (Eurogentec, France).

The protease inhibitors Aprotinin, Leupeptin, Pefablock, Sodium orthovanadate and Phosphatase Inhibitor Cocktail 3 were all obtained from Sigma-Aldrich. RPMI 1640, DPBS−/−, FCS and L-Glutamine were all purchased from Gibco (Thermo Fisher Scientific). The following chemicals were obtained from Roth: BSA, DABCO, DAPI, EDTA, Glycerol, LB, Mowiol, NaCl, Na-deoxycholate, NH_4_Cl, paraformaldehyde, SDS, Tris (hydroxymethyl)-aminoethane, Triton X-100 and Tween 20. Amikacin sulphate and Gentamicin were obtained from Sigma-Aldrich. StxB was purchased from Sigma-Aldrich (Cat. No. SML0562).

### Statistical analysis

All data in graphs are presented as mean ± standard deviation (SD) and were calculated from the results of independent experiments. Statistical testing was performed with GraphPad Prism software using data of ≥ 3 biological replicates. When appropriate, one-way analysis of variance (ANOVA), two-way ANOVA or two-tailed unpaired t-test was conducted to determine the significance of the data. Tests with a p-value < 0.05 are considered statistically significant and marked by asterisks.

## 3. Results

### 3.1 LecA preferentially binds to Gb3 species with saturated fatty acyl chains

The length and saturation level of Gb3 affects binding and subsequent trafficking of StxB [22,46], however, the influence on LecA binding is not known. First, we characterized the Gb3 species dominantly present in our cell model, the lung epithelial cell line H1299. This cell model was chosen due to the well-known impact of PA in chronic lung infections [47]. The cells were lysed in a 1:1 mixture of methanol and water. Lipids were isolated by chloroform purification and characterized by MS. Approximately 36% of the Gb3 species present were assigned to the unsaturated species C24:1. Around 32% were matched to the saturated species Gb3 C24:0 and 23% to Gb3 C16:0 (Fig. 1a). Additionally, we could detect small traces of C18:0, C18:1, C22:0 and C22:1 and hydroxylated versions of the mentioned species (9% together). To better understand the influence of Gb3 length and saturation degree on the binding of LecA, we developed a pull-down strategy with subsequent targeted liquid chromatography (LC)-MS analysis (Fig. S1). We used biotinylated LecA or StxB to stimulate H1299 cells for 20 min. Lectin-bound membrane fragments were isolated using streptavidin beads and Gb3 species pulled-down together with biotinylated LecA or StxB were assessed by MS (Fig. 1b for LecA and S2a for StxB, normalized to Gb3 C16:0; values for LecA and StxB not normalized are shown in Fig. S2b). In comparison to the tetrameric LecA, higher numbers of all detected Gb3 species were measured for the pentameric StxB, which can be explained by its binding capacity of up to 15 Gb3 lipids, while LecA exhibits only four binding sites. The normalization to the Gb3 C16:0 species allowed a comparison independent of binding avidity and demonstrated a very similar Gb3 species-preference for both lectins. Interestingly, a clear prevalence for Gb3 species with saturated fatty acyl chains (Gb3 C16:0 and C24:0) in comparison to the unsaturated Gb3 C24:1 was unraveled for both lectins, while no significant difference between C16:0 and C24:0 was detected. Importantly, the same trend was observed for the less represented Gb3 species (Figs. S2c and S2d).

**Fig. 1.**
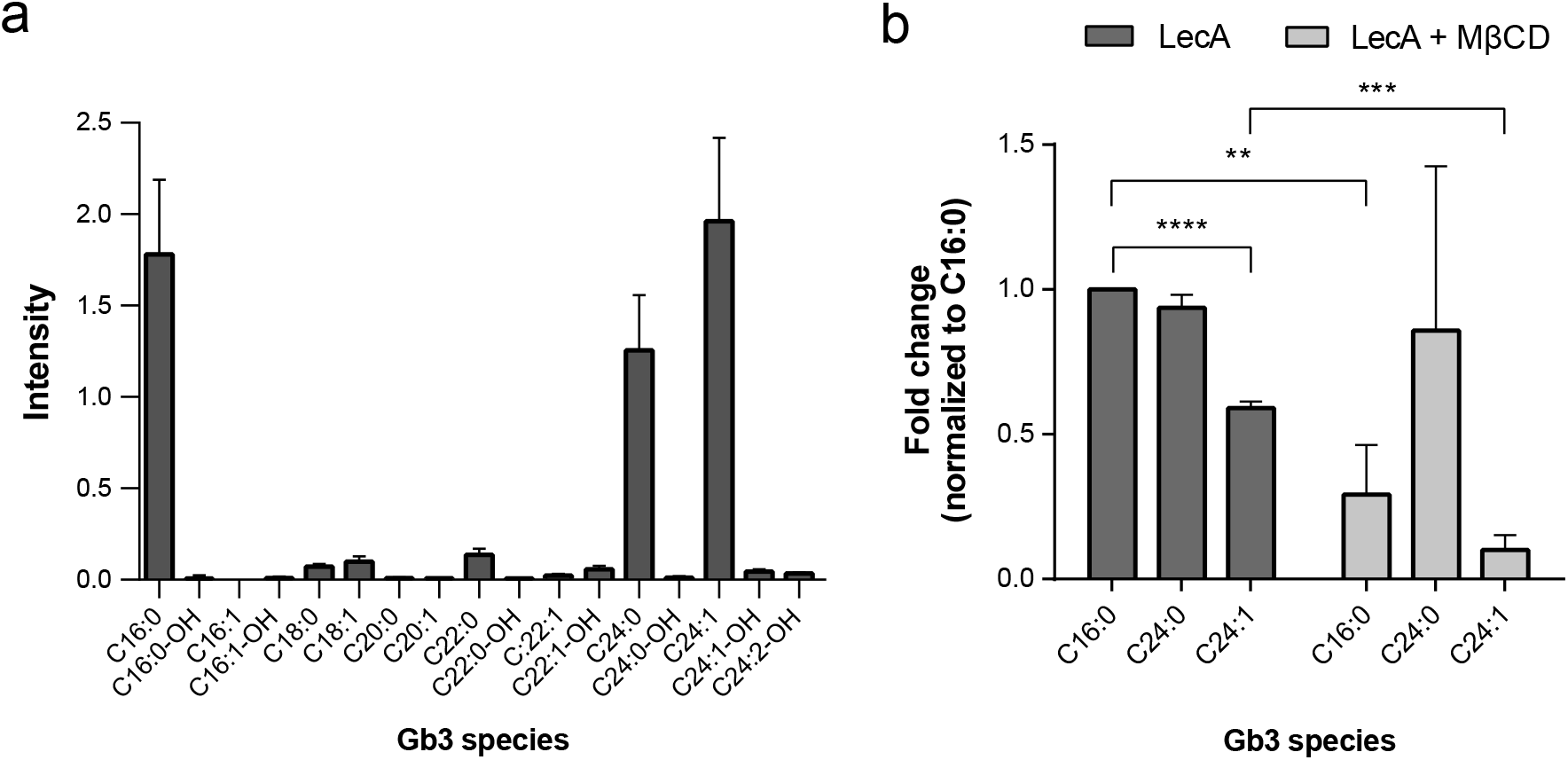
Lipid analysis by LC-MS reveals a preference of LecA for saturated Gb3 species. **a** Distribution of Gb3 species in H1299 cells. Most dominantly present Gb3 species were C16:0, C24:0 and C24:1. **b** Pulled-down Gb3 species (normalized to input values and C16:0) of LecA-biotin demonstrated a preference for saturated over unsaturated Gb3 species. Significantly less Gb3 C16:0 and C24:1 was pulled-down in MβCD-treated cells stimulated with LecA-biotin. The Gb3 species C24:0 seemed less affected by the treatment. Still, the unsaturated species C24:1 was least preferred. For clarity, only the three dominantly present species C16:0, C24:0 and C24:1 are depicted. For all panels: Bars display mean values of three biological replicates, error bars represent SD, **p<0.01, ***p<0.001, ****p<0.0001 (one-way ANOVA and Dunnett’s multiple comparisons test).

Lipid rafts are known to comprise GSLs and enriched levels of cholesterol [27]. We studied the impact of cholesterol on the binding behavior of LecA to Gb3 by depleting H1299 cells of cholesterol by MβCD treatment (10 mM, 30 min). The treatment led to a significant cholesterol reduction of ~30% in the input samples (Fig. S2e) and a strong decrease of up to 90% in the purified samples as compared to untreated H1299 cells (Fig. S2f). Subsequently, we addressed the question whether cholesterol depletion alters the Gb3 binding preference of LecA. Of note, the treatment significantly diminished binding to Gb3 overall (Fig. 1b). The strongest effect was observed for the species C16:0 and C24:1, whose binding was significantly reduced as compared to untreated cells. However, the same cannot be claimed for the saturated species C24:0 as it displayed high variability between experiments. Interestingly, the unsaturated species 24:1 of Gb3 remained the least accessible for LecA. Similar results were obtained for StxB (Fig. S2a). Our experiments, therefore, revealed a preference of the two lectins, LecA and StxB, for saturated over unsaturated Gb3 species. Overall binding to Gb3, but not the preference for saturated species was influenced by cholesterol depletion.

#### 3.2. Lipid raft proteins define the binding domain of LecA at the plasma membrane

The importance of lipids in endocytosis is not yet fully understood; however, the host proteins are integral and inevitable players of the uptake process. Membrane-associated proteins directly interact with the ligand to establish attachment points, trigger signaling pathways, scaffold the binding domain, or aid in membrane bending and scission processes. We aimed at identifying novel protein interaction partners of LecA within the Gb3 membrane domain following stimulation of H1299 cells with LecA-biotin for 5 and 15 min. The lysed membrane fragments were processed as described before (section 3.1) and pulled-down proteins were on-bead trypsin-digested prior to MS analysis. The protein hits included various cytoskeleton components and cytoskeleton-regulating proteins (Table S2). Already after 5 min of LecA incubation, vimentin was highly enriched in the LecA-treated sample and after 15 min of stimulation actin, tubulin, myosin-9 and the small GTPase Rac1 were detected. Additionally, the GPI-anchored protein CD59 and the Src kinase Yes were identified. Together with the scaffolding protein flotillin-1, which accumulated after 15 min, these proteins represent well-reported lipid raft components [48].

To characterize the identified hits in more detail, we preformed confocal microscopy, co-immunoprecipitation and immunoblotting experiments (Figs. 2 and 3). Fluorescently labeled LecA co-localized with flotillin-1 (Fig. 2a), flotillin-2 (Fig. S3a) and CD59 (Fig. 3a) in immunofluorescence studies. The fluorescence signal overlapped predominantly at the plasma membrane (see white arrows in Figs. 2a, 3a and S3a) but also in vesicular structures and perinuclear region (marked by asterisks in Figs. 2a, 3a and S3a). In contrast to CD59, flotillins were detected almost exclusively in the perinuclear region in untreated control cells but re-localized to the plasma membrane upon LecA treatment (Figs. 2a and S3a). Co-localization was quantified and revealed an increasing overlap between LecA and flotillins or CD59 over time (Figs. 2b, 3b and S3b). Additionally, we performed a pull-down of LecA-biotin and confirmed the presence of the two flotillins and CD59 in the LecA plasma membrane domain biochemically by immunoblotting (Figs. 2c, 3c and S3c). Of note, the pull-downs suggest a time-dependent recruitment of the proteins to the cell surface indicated by increasing protein levels over time. Both, flotillin-1 and CD59 could be detected together with LecA in the eluates of the corresponding co-immunoprecipitation (co-IP) of LecA-treated H1299 cell lysates (Figs. 2d and 3d). The co-precipitation was glycan-dependent as PPMP treatment, an agent that inhibits the synthesis of glycosylceramide-based GSLs [41], strongly diminished the interaction of LecA with flotillin-1 or CD59. However, we neither witnessed a direct binding of LecA to flotillin-1 nor CD59 as shown by immunoblots of IP eluates, which were incubated with LecA-biotin and fluorescent streptavidin (Fig. S4). We, therefore, conclude that LecA binding to Gb3 induces the recruitment of flotillins and CD59 into LecA/Gb3 membrane domains.

**Fig. 2.**
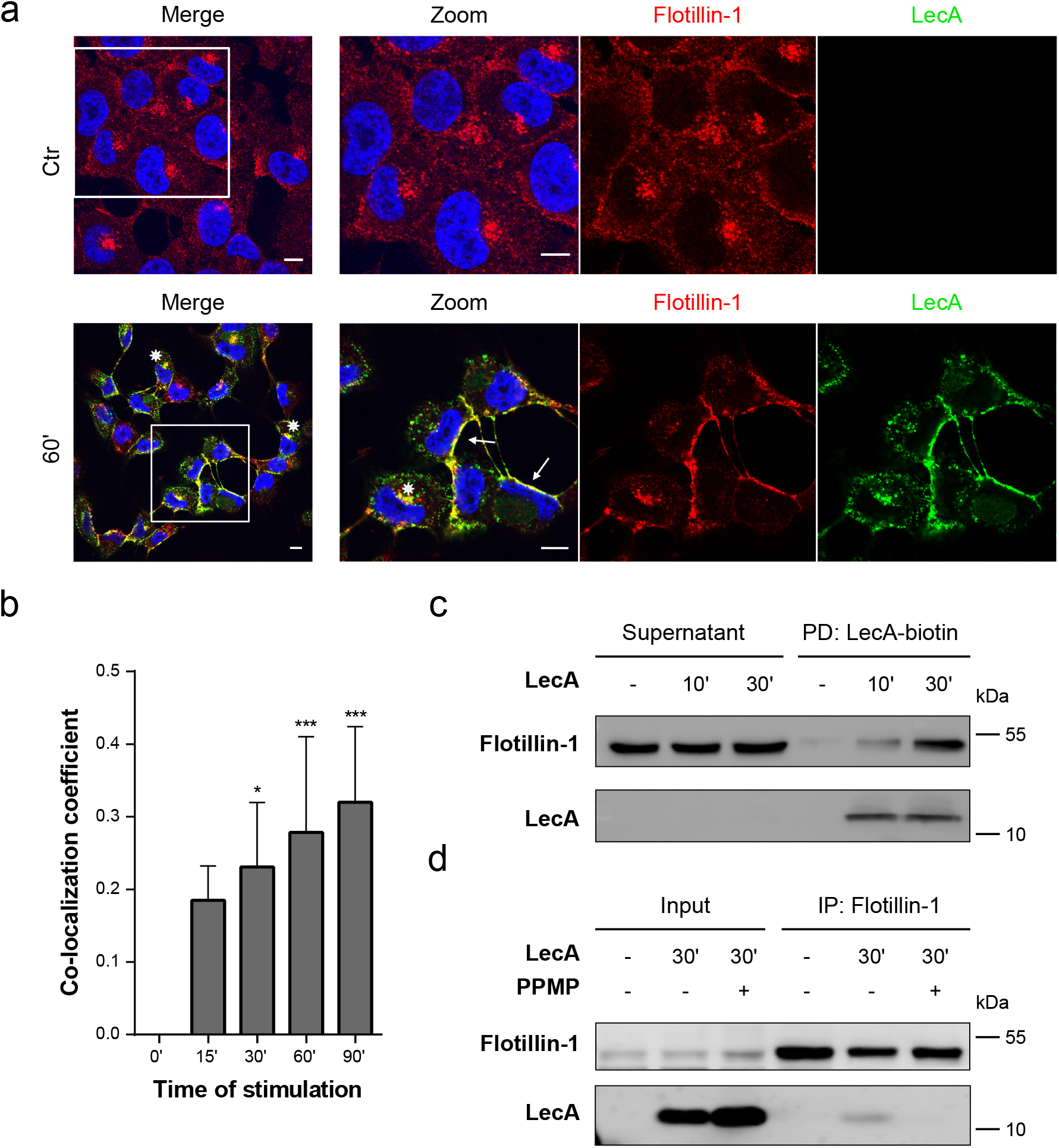
Flotillins are recruited to the plasma membrane upon LecA stimulation. **a** Fluorescence co-localization studies of flotillin-1 (red) and LecA (green) after 60 min of lectin stimulation. Nuclei were counterstained by DAPI. Framed areas were magnified. White arrows point at co-localization events at the plasma membrane, asterisks at perinuclear co-localization. Scale bar: 10 μm. **b** Mander’s co-localization coefficient quantified between the fluorescence signals of flotillin-1 and LecA in comparison to time point 0. Bars display mean values of at least three biological replicates, error bars represent SD, *p<0.05, ***p<0.001 (one-way ANOVA and Dunnett’s multiple comparisons test). **c** Pull-down of LecA-biotin resulted in a time-dependent enrichment of flotillin-1. **d** LecA was co-immunoprecipitated with flotillin-1 after 30 min of LecA treatment, the binding and precipitation was inhibited by PPMP treatment.

**Fig. 3.**
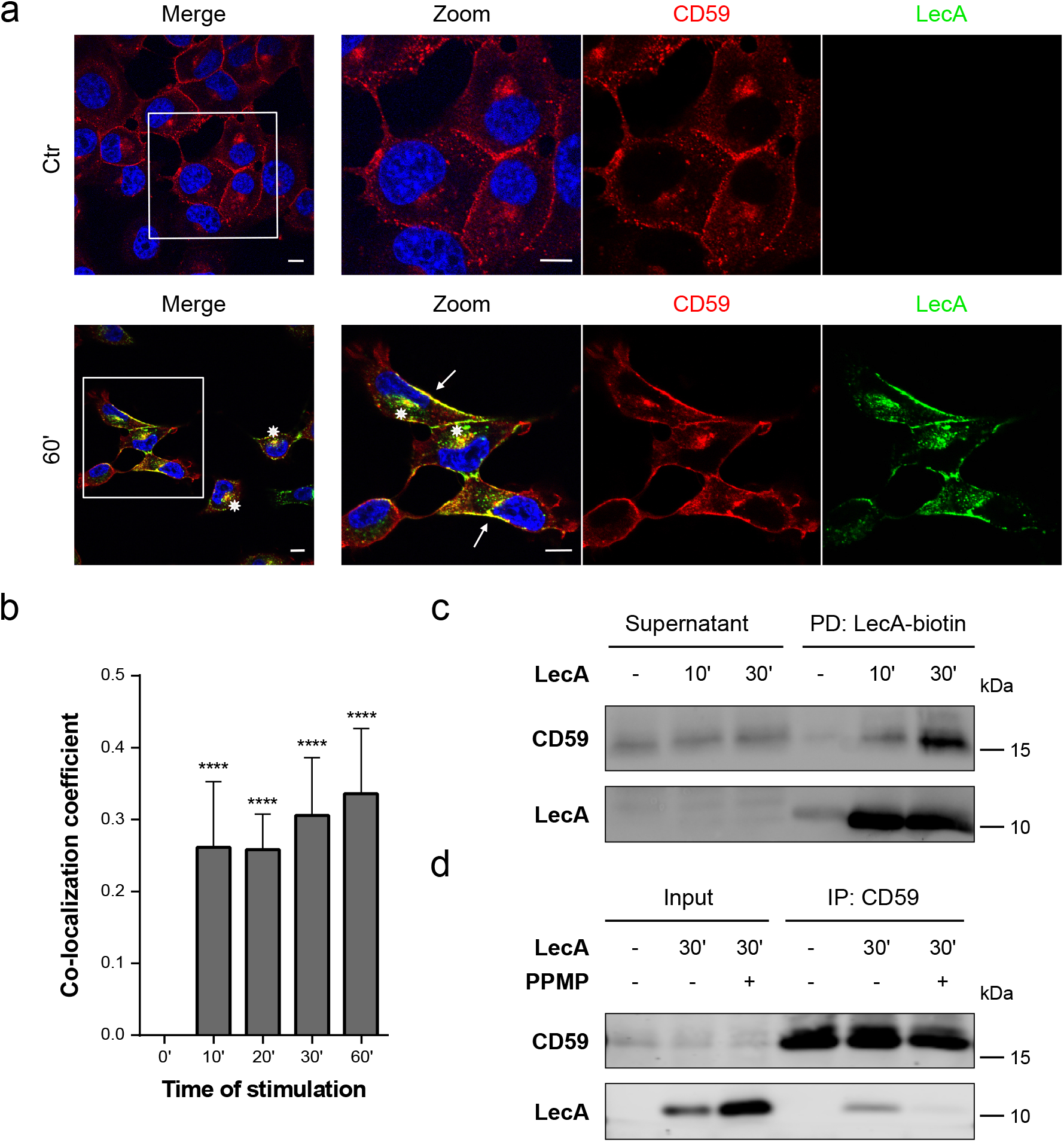
The LecA plasma membrane domain includes CD59 proteins. **a** Fluorescence co-localization studies of CD59 (red) and LecA (green) after 60 min of lectin stimulation. Nuclei were counterstained by DAPI. Framed areas were magnified. White arrows point at co-localization events at the plasma membrane, asterisks at perinuclear co-localization. Scale bar: 10 μm. **b** Signal overlay of LecA and CD59 is displayed by Mander’s co-localization coefficient quantified in comparison to time point 0. Bars display mean values of at least four biological replicates, error bars represent SD, ****p<0.0001 (one-way ANOVA and Dunnett’s multiple comparison tests). **c** CD59 was validated as component of the LecA-binding domain by pull-down of LecA-biotin. **d** Immunoprecipitation of CD59 co-precipitated LecA after 30 min of stimulation. PPMP treatment in order to deplete cells in glycosylceramide-based GSLs, such as Gb3, inhibited the precipitation of LecA.

#### 3.3 LecA activates Src kinases and Rac-1

We recently demonstrated that the adaptor protein CrkII is activated upon stimulation with LecA in H1299 cells [17]. Src family kinases were identified as a crucial upstream factor of LecA-induced CrkII activation. Since we detected the Src kinase Yes in the protein MS analysis, we further analyzed the role of Src family kinases in processes related to LecA binding. Using immunoblot analysis, we confirmed the LecA-mediated increase in phosphorylation of Src kinases at Tyr^416^ in a time-dependent manner (Figs. 4a and b, also compare with [17]). We also validated the presence of Src in the LecA plasma membrane domain by pull-down of LecA-biotin (Fig. 4c). We performed Rac1 G-LISA to validate the role of Rac1, an important GTPase found in our MS hits (Fig. 4d). The small GTPase was activated upon LecA stimulation, however, great variability between the biological replicates was observed. Potentially, this was owed to the short lifetime of Rac1 activation.

**Fig. 4.**
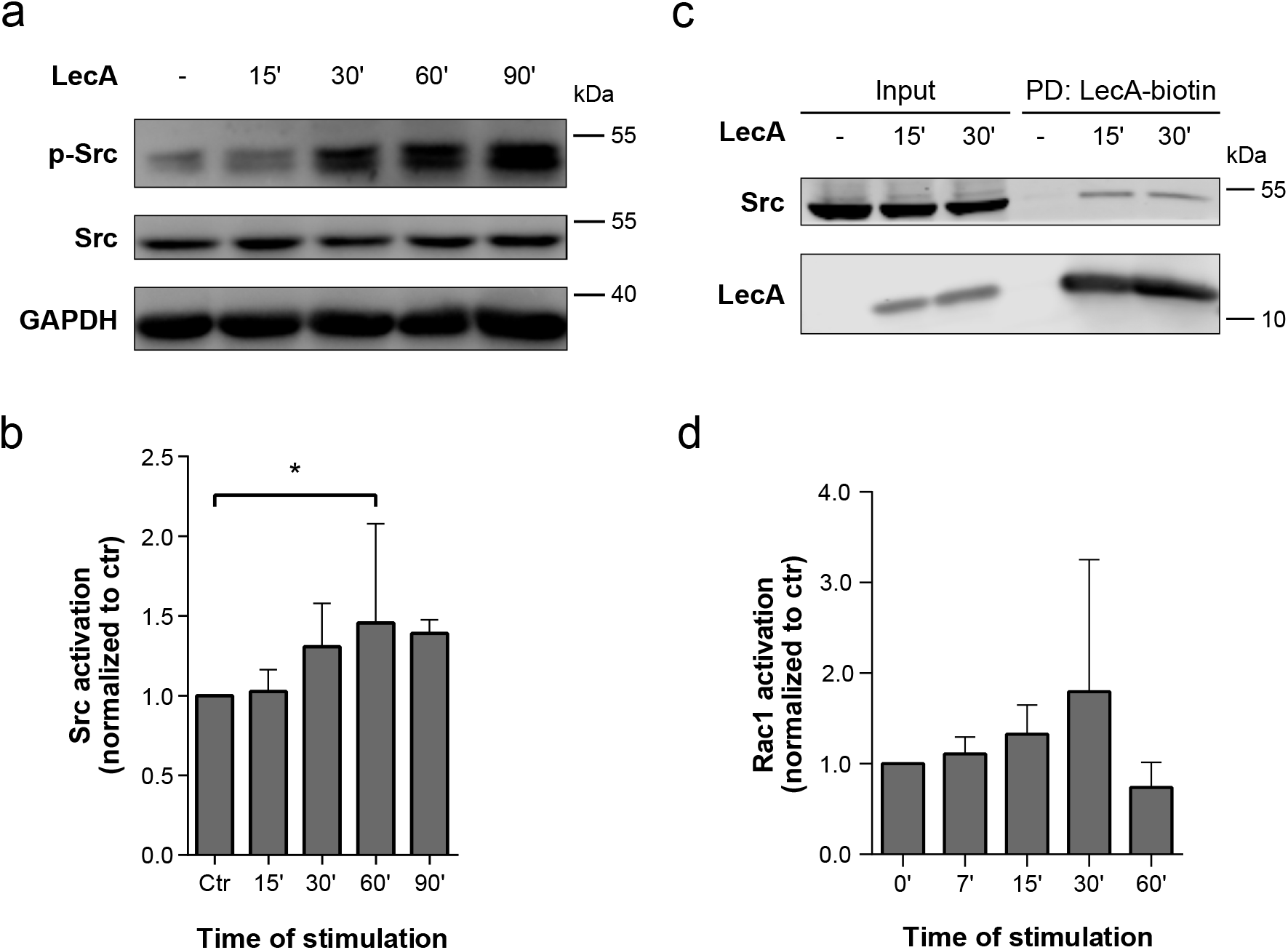
Src family kinases and Rac1 are activated upon LecA stimulation. **a** Src family kinases are phosphorylated in H1299 cells upon LecA treatment as demonstrated by immunoblot analysis. **b** Src activation was quantified by the ratio between phospho-Src (Tyr^416^) and Src and normalized to control levels. Bars display mean values of at least four biological replicates, error bars represent SD, *p<0.05 (one-way ANOVA and Dunnett’s multiple comparisons test). **c** The Src family kinase was pulled-down together with LecA after 15 and 30 min of treatment. **d** G-LISA of Rac1 demonstrated a non-significant activation of the small G protein Rac1 upon LecA stimulation. Values were normalized to control levels. Bars display mean values of three biological replicates, error bars represent SD.

Taken together, our data demonstrate that LecA assembles a flotillin/CD59-enriched plasma membrane domain and induces the recruitment of Src kinases and the small GTPase Rac1 for the activation of cellular mechanisms. These factors in combination with the PI3-kinase signaling cascade represent critical players in the process of actin reorganization [49–51].

#### 3.4 PIP_3_ clusters are induced upon LecA treatment

Phosphatidylinositols (PIPs) naturally bear a long saturated C18 acyl chain and, therefore, reach across bilayer enabling an interaction with long chains of lipids in the extracellular leaflet [52,53]. To investigate a potential coupling mechanism between the extracellular and the intracellular leaflet of the host cell plasma membrane, we studied the influence of LecA-treatment on phosphatidylinositol (4,5)- bisphosphate (PIP_2_) phosphorylation in H1299 cells. The PH domain of the PI3-kinase downstream signaling molecule Protein kinase B (Akt) can sense PIP_3_ and is commonly used as a GFP-fusion protein to study PIP_3_ dynamics in living cells [54,55]. We transfected H1299 cells with PH-Akt-GFP and traced GFP signal for 60 min following fluorescent LecA-treatment. Prior to stimulation with LecA, cells displayed an evenly distributed GFP signal in the cytosol and at the plasma membrane (Fig. 5, upper panel). Upon stimulation with LecA, PH-Akt-GFP clusters appeared at the LecA binding site indicating phosphorylation of PIP_2_ through activated PI3-kinases and clustering of PIP_3_ in the intracellular leaflet of the plasma membrane (Fig. 5, lower panels). The first PIP_3_ clusters appeared within 2-3 minutes post treatment as depicted in the animation (Online Resource 1). Over the whole time-course, the PIP_3_ clusters displayed a dynamic character with some clusters disappearing once LecA was endocytosed (orange arrows in Fig. 5).

**Fig. 5.**
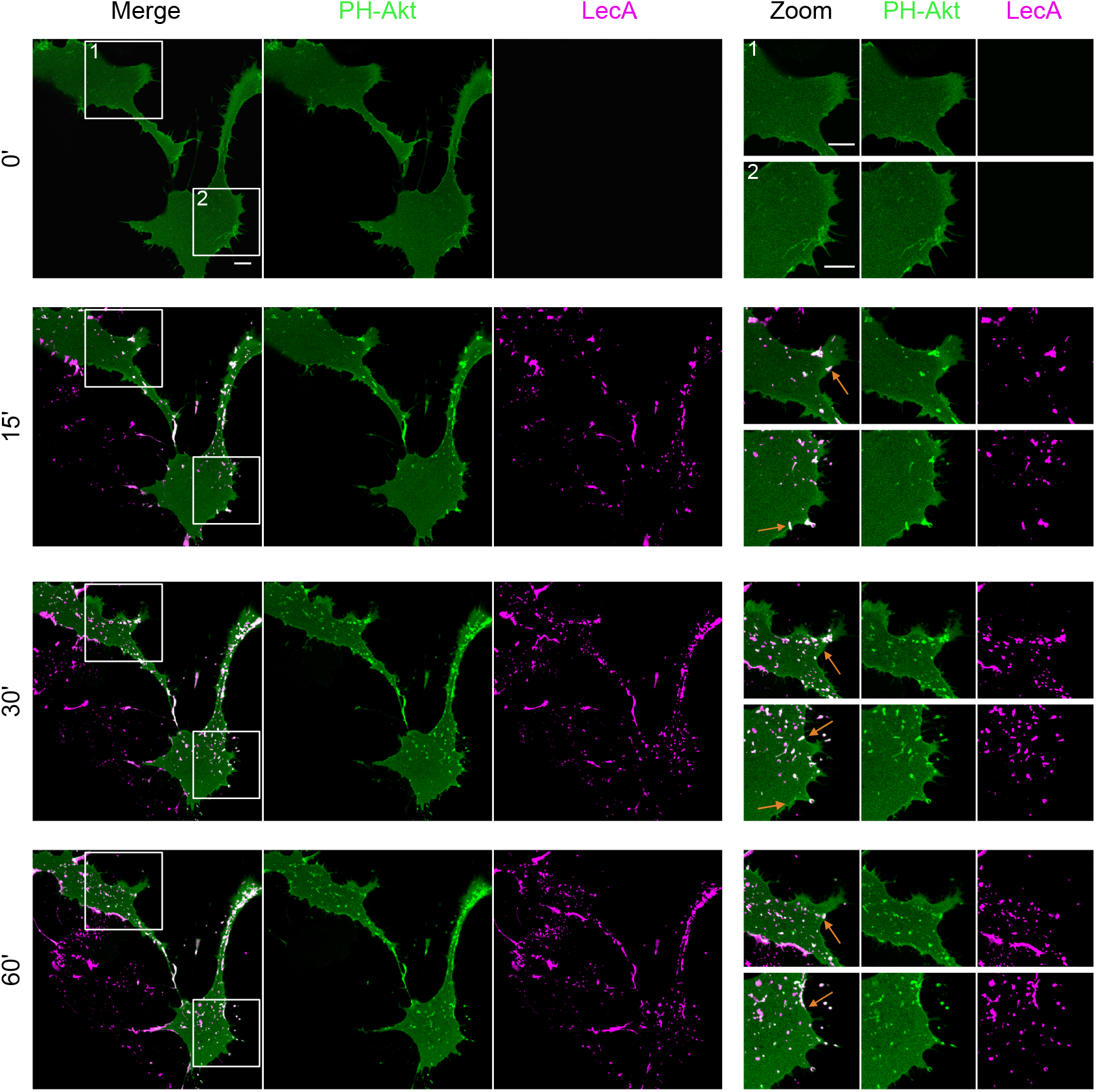
LecA induces clustering of PIP_3_ in H1299 cells. **a** Left panel: Time-lapse images of PH-Akt-GFP expressing H1299 cells exposed to fluorescent LecA over 60 min. Right panel: Magnifications highlight dynamic co-localizing events between LecA and PH-Akt-GFP correlating with LecA endocytosis (marked by orange arrows). Framed areas were magnified. Scale bar: 10 μm.

These observations encouraged us to study a potential link between the PI3-kinase and flotillins by co-transfection of H1299 cells with PH-Akt-GFP and flotillin-1-mCherry (Fig. 6). Strikingly, a clear co-localization of all three proteins (PH-Akt, flotillin-1 and LecA) was observed after 30 min of stimulation (Fig. 6a). As described before, flotillin-1 was recruited to the plasma membrane and accumulated at LecA nucleation domains. Addition of Wortmannin, a potent inhibitor of PI3-kinase activity [56], disrupted the co-localization of PH-Akt-GFP and LecA (Fig. 6b) and also fewer PH-Akt-GFP clusters were detected. Interestingly, this reduced the co-localization of flotillin-1-mCherry and LecA as well (Fig. 6c). Moreover, Wortmannin-treatment abrogated recruitment of flotillin-1 to the plasma membrane. Similar results were obtained for a co-transfection of PH-Akt-GFP and flotillin-2 (Fig. S5). Taken together, we propose that the LecA-induced recruitment of flotillins to the plasma membrane depends on PI3-kinase activity leading to phosphorylation of PIP_2_. The induced processes are important for efficient internalization of LecA.

**Fig. 6.**
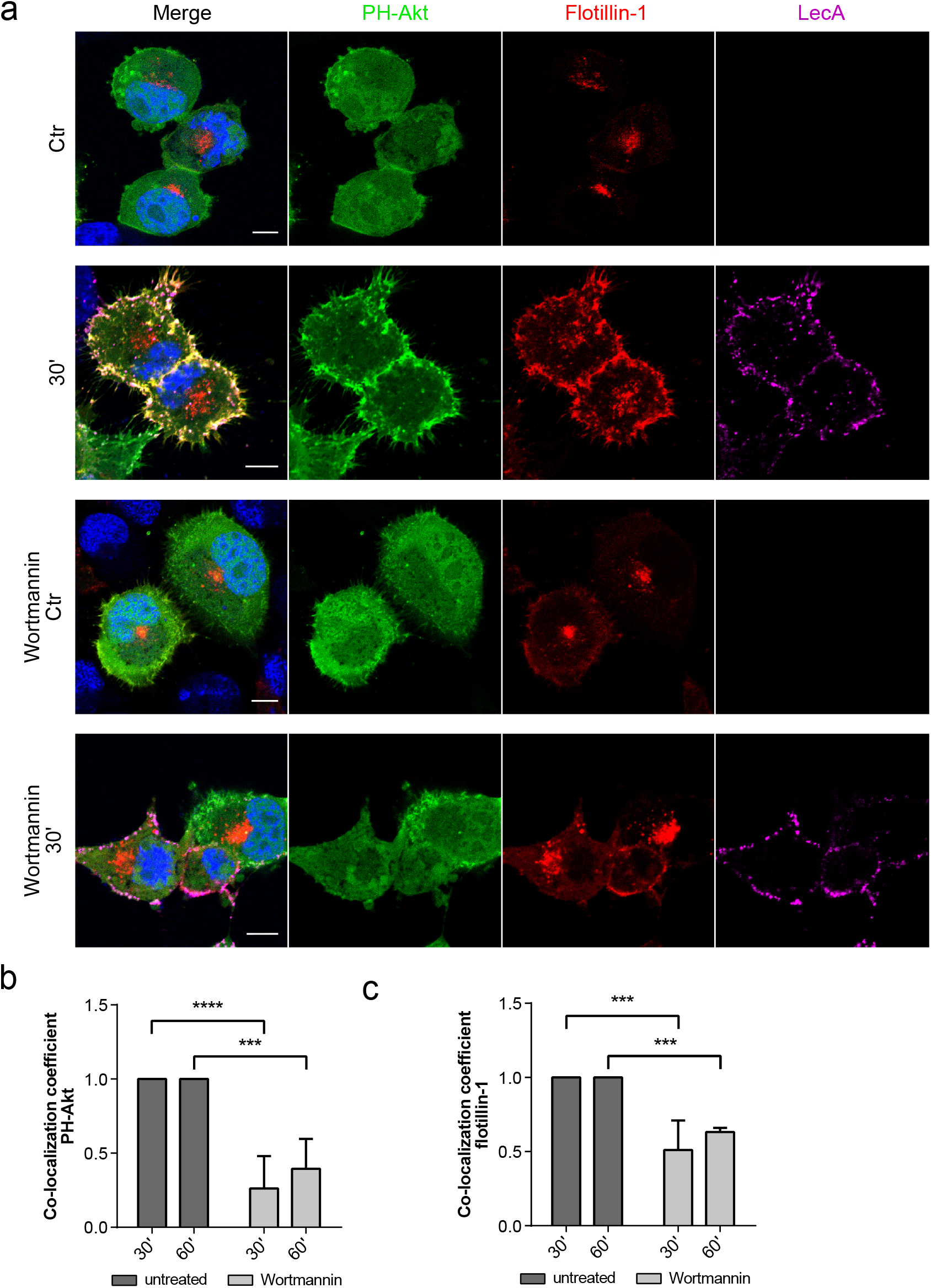
PI3-kinase inhibition partially prevents PIP_3_ clustering and recruitment of flotillins upon LecA stimulation. **a** Confocal microscopy images of PH-Akt-GFP and flotillin-1-mCherry expressing H1299 cells exposed to fluorescent LecA. Lower two panels: Cells were pre-treated with 100 nM Wortmannin to inhibit PI3-kinase activity. Scale bar: 10 μm. **b** Co-localization of PH-Akt-GFP and LecA is depicted as fold change of Mander’s co-localization coefficient normalized to the untreated conditions. **c** Fold change of Mander’s co-localization coefficient quantified between the fluorescence signals of flotillin-1-mCherry and LecA in comparison to the untreated conditions. For all panels: Bars display mean values of three biological replicates, error bars represent SD **p<0.01, ***p<0.001, ****p<0.0001 (two-way ANOVA and Tukey’s multiple comparisons tests).

#### 3.5 CD59 and flotillins strongly affect the invasiveness of *P. aeruginosa* into H1299 cells

LecA is crucial for the pathogenicity of PA [5–8]. Here, we identified the proteins CD59, flotillin-1 and −2, the Src family kinases, Rac1 and PIP_3_ as components of the plasma membrane domain for LecA-induced signaling and entry. Src kinases, small GTPases and PIPs are host cell factors with an established role in PA infection [16,57,58]. CD59 and flotillins, however, have not been studied in this context yet and we, therefore, aimed to understand the impact of these newly identified proteins for host cell invasion of PA. The invasion efficiency of PAO1 was analyzed with respect to the presence of flotillins and CD59 in H1299 cells (Fig. 7). To do so, we first created a CRISPR-Cas9 knockout model of *FLOT1* (Δ*FLOT1*) in H1299 cells (Fig. S6). To negate the effects of flotillin-2, we additionally silenced *FLOT2* expression by transfection of Δ*FLOT1* cells with siRNA (Fig. 7a). Here, the interdependency of the two flotillins could be highlighted in an immunoblot of WT and Δ*FLOT1* lysates additionally subjected to flotillin-2 silencing: flotillin-1 levels decreased in WT cells transfected with flotillin-2 siRNA, while less flotillin-2 was detected in Δ*FLOT1* cells as compared to H1299 WT. Additionally and more importantly, the knockdown of flotillin-2 by siRNA was more efficient in Δ*FLOT1* cells. We were therefore able to conduct experiments in H1299 cells almost entirely depleted of flotillins. Similarly, expression of CD59 was silenced by *CD59* siRNA transfection of H1299 WT cells (Fig. 7b).

**Fig. 7.**
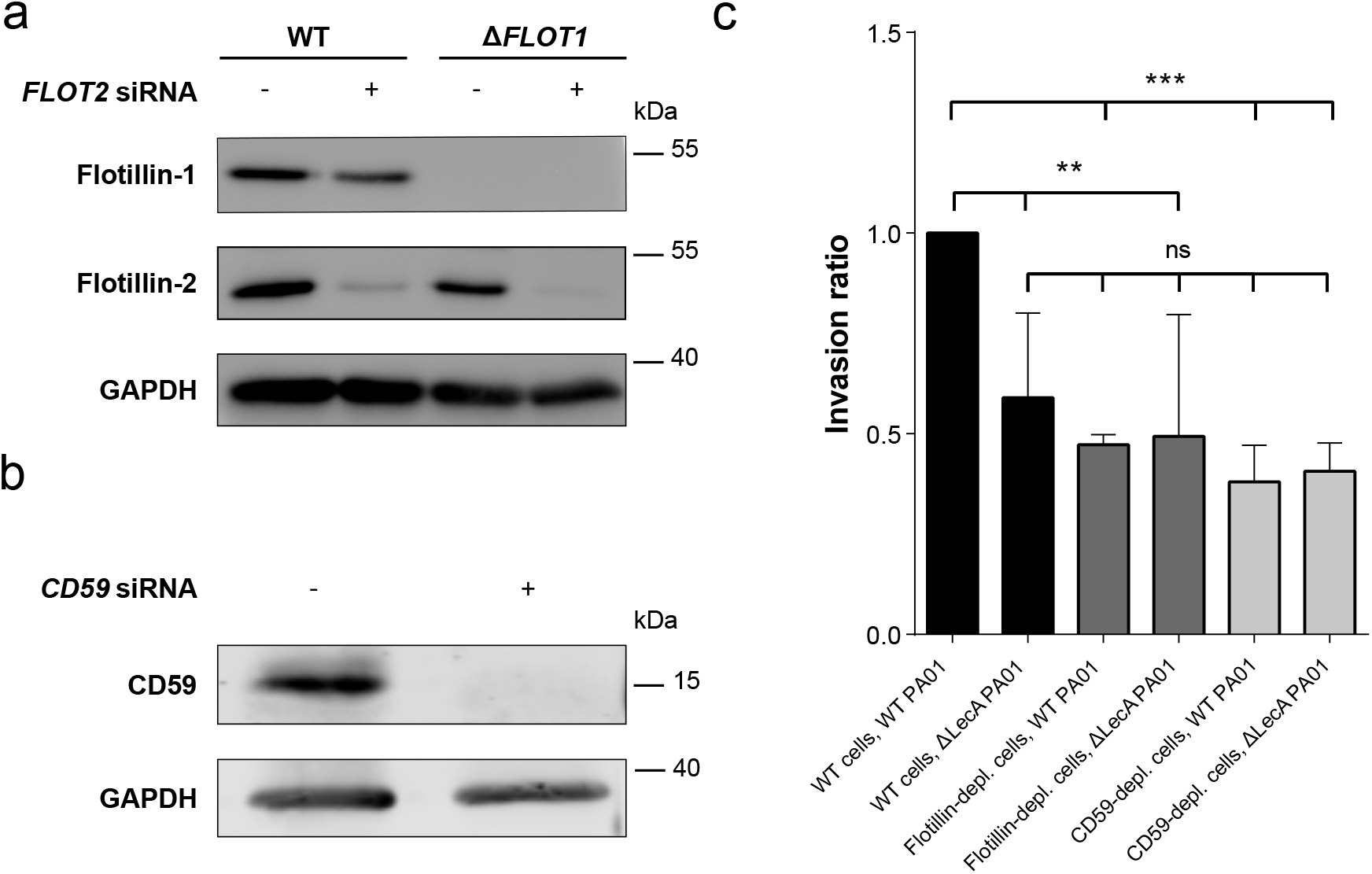
Flotillins and CD59 promote cellular invasion of *P. aeruginosa* in conjunction with LecA. **a** CRISPR-Cas9 knockout of *FLOT1* and siRNA transfection of *FLOT2* depleted H1299 cells almost completely of flotillins. Note, that protein levels of flotillins are interdependent. **b** CD59 was efficiently knocked-down by siRNA transfection of WT H1299 cells. **c** Invasion assay of WT, flotillin- and CD59-depleted H1299 cells using WT PAO1 and Δ*LecA* mutant PAO1 strains demonstrated the impact of flotillins and CD59 for PAO1 invasion. Invasion ratio was normalized to WT cells and WT PAO1. Bars display mean values of at least three biological replicates, error bars represent SD, **P<0.01. ***P<0.001 (one-way ANOVA and Dunnett’s multiple comparisons test).

Strikingly, the experiments demonstrated a reduction in PA invasion into H1299 cells by more than 50% or 60% in cells depleted of flotillins or CD59, respectively (Fig. 7c). These observations clearly point towards a promotional role of the two proteins for PA invasion. Similarly, infection with the Δ*LecA* PAO1 mutant strain reduced the invasion efficiency into H1299 WT host cells [8] to comparable levels of CD59- or flotillin-depleted cells. Conversely, infection of CD59- or flotillin-depleted cells with Δ*LecA* PAO1 did not further decrease the invasion efficiency, which suggests that LecA, CD59 and flotillins act in the same pathway. Our approach furthermore demonstrates the successful transfer of results obtained by studying a single bacterial virulence factor, namely LecA, to the complete bacterium - a process that facilitates the identification of key players of infection.

### 4. Discussion

Membrane processes related to the uptake of PA into host cells are only beginning to be elucidated, as this bacterium is mostly considered as an opportunistic extracellular pathogen [59]. However, at certain sites of colonization, for example in the lungs, invasive strains of PA cause severe and chronic infections. So far, PIPs [58], the Src family kinases [57,60], the actin cytoskeleton and the microtubule network [8,61,62] are known host factors necessary for the internalization of PA. Here, we demonstrate that the flotillin family and the GPI-anchored protein CD59 (one of the cargo proteins of flotillin-assisted endocytosis) play a significant role for PAO1 invasion into lung epithelial cells. Flotillins are evolutionary conserved and ubiquitously expressed proteins with particularly high expression levels in heart, brain and lungs [63]. Nevertheless, their cellular functions are incompletely understood and their role in endocytosis is controversial. On the one hand, Glebov and colleagues investigated the uptake of CD59 and GM1, the receptor for cholera toxin, and demonstrated a requirement for flotillins and dynamin but not for clathrin in endocytosis of these cargos [31]. On the other hand, several studies suggest that flotillins might work hand-in-hand with clathrin-mediated endocytosis by clustering of cargos prior to their internalization via clathrin [39,64–66]. Furthermore, flotillins coordinate cargo sorting, recycling and trafficking of several toxins without an influence on their uptake *per se* [67–69]. Similarly, little is known about the interplay between flotillins and pathogens. Korhonen and colleagues demonstrated a reduced intracellular growth of *Chlamydia pneumoniae* in the absence of flotillin-1 [70]. In addition, the invasion of erythrocytes by the parasite *Plasmodium falciparum* is dependent on lipid rafts and the recruitment of both flotillin proteins and CD59 to the parasitophorous vacuole [71–73]. A recent study strengthens the available data by highlighting a critical role of flotillins in *Anaplasma phagocytophilum* infection [74].

The intricacy of bacteria and the host plasma membrane complicates research on a molecular level. By reducing the complexity of the system and analyzing the interaction of a single bacterial factor, namely LecA, with the host cell plasma membrane we gained insights into the key players of binding and uptake processes and determined important parameters for the invasiveness of *P. aeruginosa* strains. Interaction partners of LecA were identified by purification of proteins directly or indirectly bound by LecA-biotin using proteomic analysis. Strikingly, many detected proteins are known lipid raft components including the GPI-anchored protein CD59, the flotillin proteins and the Src family kinase Yes [75–77]. The complement regulation factor CD59 was already functionally linked to Src family kinases in studies demonstrating the recruitment and activation of trimeric G proteins and the Src kinase Lyn to CD59 clusters in the plasma membrane [75,78]. Additionally, CD59 is a known cargo of flotillin-assisted endocytosis [31] and highlighted in several studies as an interaction partner of flotillins [79,80].

Moreover, both flotillins can be tyrosine-phosphorylated by members of the Src family kinases and closely associate at the plasma membrane [77,81]. As we previously demonstrated [17] and verified here, Src family kinases are activated and phosphorylated at Tyr^416^ upon LecA treatment of H1299 cells. The activation of Src kinases might eventually culminate in cytoskeletal reorganization, a prerequisite for the uptake of lectins and whole bacteria. Furthermore, we detected known endocytic players of lipid rafts in our MS screen. These include the small G protein Rac1 and cytoskeletal components like vimentin and myosin 9, of which the latter, interestingly, is regulated by flotillins [51]. Conversely, all factors might work together to orchestrate the uptake of LecA and PAO1.

Assembly of transmembrane receptors and scaffolding proteins into specialized membrane domains is as important as the clustering of lipids on both membrane leaflets for enabling interactions with recruited proteins and signal transduction. To further characterize the LecA-induced plasma membrane domain, we analyzed the Gb3 species pulled-down together with LecA-biotin. The three most abundant Gb3 species present in H1299 cells are C16:0, C24:0 and C24:1 as determined by lipid MS. These results agree with data presented in [82], where the composition of Gb3 species in several cell lines is summarized. Strikingly, the saturated Gb3 species C16:0 and C24:0 pulled-down by LecA were enriched roughly by a factor of 1.7 in comparison to the unsaturated species C24:1. The saturation level of Gb3 ceramide tails has been shown to drastically affect lipid bilayer phases [83]. Therefore, it is unsurprising that several studies describe the critical impact of acyl chain length and degree of saturation on binding and trafficking of protein toxins [21,26,84,85]. The detected preference of LecA towards saturated Gb3 species in our MS analysis suggests preferential binding to rather tightly packed membrane domains, a characteristic feature of lipid rafts. Gb3 species with little abundance in the plasma membrane in our study, e.g. hydroxylated Gb3 species, were reported to play a crucial role in the scission of StxB-induced membrane tubules [86].

Several studies investigated the binding of StxB to its receptor Gb3 in synthetic model membranes with diverse results [22,84,85]. Interestingly, it was shown for StxB and simian virus 40 that successful trafficking to the endoplasmic reticulum (ER) and induction of toxicity requires binding to long saturated GSLs [87–89], while cholera toxin only sorted efficiently from the plasma membrane to the Golgi network and ER by interaction with unsaturated ceramide chains of its receptor, the GSL GM1 [69]. This provides further evidence that subtle changes in the lipid bilayer influence carbohydrate exposure and consequently allow, weaken or restrict lectin binding. Furthermore, the surrounding membrane environment is critical for Gb3 receptor function [90,91]. Depletion of cholesterol by MβCD significantly decreased levels of isolated Gb3 C16:0 and C24:1 species. The amount of measured Gb3 C24:0 varied between the three replicates but did not change significantly. The predilection of LecA-biotin towards the saturated Gb3 species was, however, maintained.

Of note, LecA and StxB both bind to Gb3, but partially localize to different membrane domains [23] and exhibit distinct trafficking routes [92]. In our analysis, no clear difference in binding behavior to Gb3 species of LecA in comparison to StxB was observed. Therefore, we suggest that the fates of the endocytosed lectins are rather determined by the set of cellular interaction partners responsible for downstream signal transduction and cargo sorting.

The binding of LecA to Gb3 generates a signal at the extracellular leaflet but so far no transmembrane-spanning protein linking Gb3 with the intracellular membrane leaflet is known. How is the signal then communicated to flotillins and Src kinases? A long-standing, popular hypothesis suggests a role for fatty acids in signal transduction [19,93]. Pinto and colleagues studied the influence of ceramide (i.e. the backbone of Gb3) structure and acyl chain length and demonstrated that long-chain ceramides (C24) induce strong alterations in the bilayer, suggesting the formation of interdigitating phases [94]. Further, GPI-anchored proteins with long saturated acyl chains can interdigitate and connect the membrane leaflets given that one of the two leaflets is immobilized [20]. A good example of transbilayer coupling through GPI-anchored proteins might be the prion protein (PrP). PrP clustering in the extracellular leaflet was suggested to influence flotillins at the intracellular leaflet through interaction with flotillin myristoyl- and palmitoyl-residues [38]. Recently, PIPs were proposed to participate in transbilayer coupling [53]. Since PIPs naturally bear a long saturated C18 acyl chain, they reach across the bilayer and interact with long fatty acyl chains of lipids located in the extracellular leaflet. Here, we demonstrate an induction of PIP_3_ clustering upon LecA stimulation of H1299 cells. Cluster formation was crucial for flotillin recruitment to the plasma membrane. Co-clustering of long, saturated Gb3 species and the GPI-anchored protein CD59 with PIPs might therefore enable the communication across the plasma membrane, inducing intracellular processes that culminate in endocytosis. Flotillins functionally scaffold and enhance signal transduction [32,48]. Since internalization of PA is known to require PI3-kinase and Src family kinase activity [57,58,60], the induction of these processes by LecA primes the cellular plasma membrane for PA uptake (Fig. S7, potential model).

### 5. Conclusion

In this study, we characterize the host cell plasma membrane domain to which the *P. aeruginosa* virulence factor LecA binds selectively. This membrane domain is composed of saturated Gb3 species along with the GPI-anchored protein CD59. Upon LecA binding and clustering of Gb3, intracellular flotillins are recruited and scaffold active signaling platforms including Src family kinases. Flotillin recruitment is dependent on active PI3-kinases. The lectin-induced signal may therefore be transmitted from the extra-to the intracellular leaflet by transbilayer coupling between long fatty acyl chains of Gb3 or CD59 and PIP_3_. In addition, we unravel a crucial role of flotillins and CD59 in the invasion of PAO1 into lung epithelial cells. We, therefore, propose a model in which LecA prepares and reorganizes the plasma membrane for the entry of *P. aeruginosa*.

## Declarations

### Ethics approval

Not applicable

### Availability of data and material

The datasets generated and/or analyzed during the current study are available from the corresponding author on reasonable request.

### Code availability

Not applicable

### Author contributions

Conception and design of the research: AB, JM, TE, BK, WR; Funding acquisition: BK, WR; Investigation: AB, SA, SL, MS, AL, DH; Methodology: SL, MS, DF, AVM; Supervision: JM, TE, BK, WR; Writing – original draft: AB; Writing – review & editing: SA, SL, JM, TE, WR. All authors agreed on the final version of the manuscript.

## Abbreviations

emPAI: Exponentially Modified Protein Abundance Index
Δ*FLOT1*: CRISPR-Cas9 knockout model of *FLOT1*
DPPS: Dulbecco’s phosphate-buffered saline
Gb3: globotriaosylceramide
GPI: glycosylphosphatidylinositol
GSL: glycosphingolipid
IP: immunoprecipitation
LC: liquid chromatography
MS: mass spectrometry
MβCD: methyl-beta-cyclodextrin
PA, PAO1: *Pseudomonas aeruginosa*
PI3-kinase: phosphoinositide 3-kinase
PIP_2_: phosphatidylinositol (4,5)-bisphosphate
PIP_3_: phosphatidylinositol (3,4,5)-trisphosphate
PIPs: phosphatidylinositols
PPMP: D-threo-l-phenyl-2-palmitoylarmino-3-morpholino-l-propanol
PrP: prion protein
RIPA: radio-immunoprecipitation assay
RPMI: Roswell Park Memorial Institute
RT: room temperature
SD: standard deviation
sgRNA: single guide RNAs
StxB: Shiga toxin B-subunit
WT: wild type

