## Supplementary figures for "The Gb3-enriched CD59/flotillin plasma membrane domain regulates host cell invasion by *Pseudomonas aeruginosa*"

### Supplementary figures and figure legends

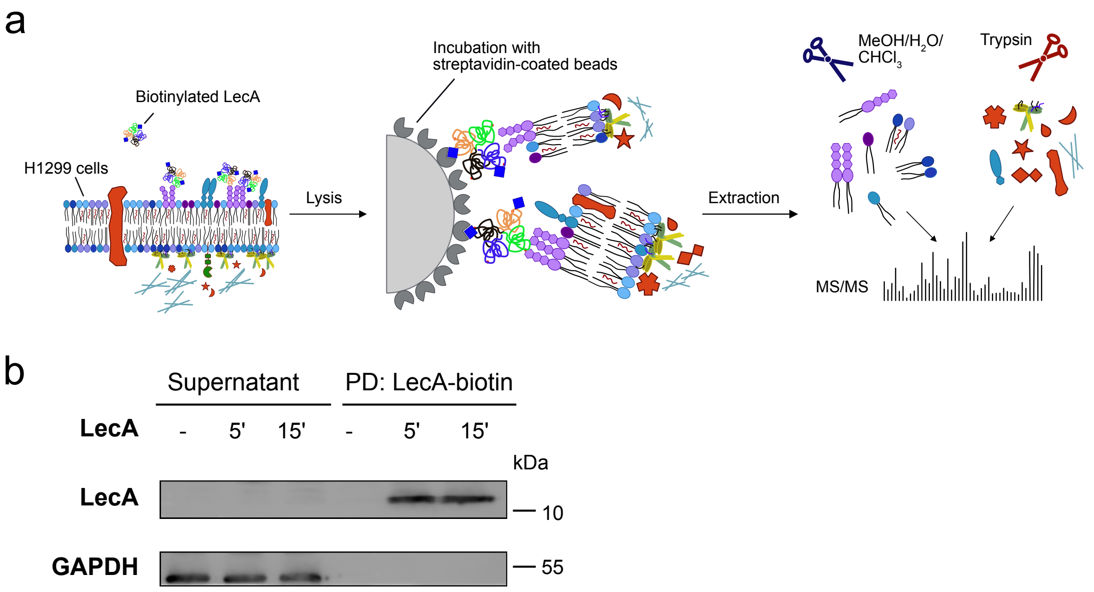


**Fig. S1** Experimental design of the study. **a** H1299 cells were incubated with biotinylated LecA for 5 and 15 min and lysed with a medium-harsh lysis buffer. The LecA-bound membrane fragments were isolated by streptavidin-coated beads and subsequently eluted using two different techniques. Depending on the target of interest, lipids or proteins were extracted from beads by methanol/water/chloroform or trypsin, respectively. After separation and purification, samples were analyzed by MS. **b** Pull-down efficiency tested by immunoblot analysis after several rounds of optimization.

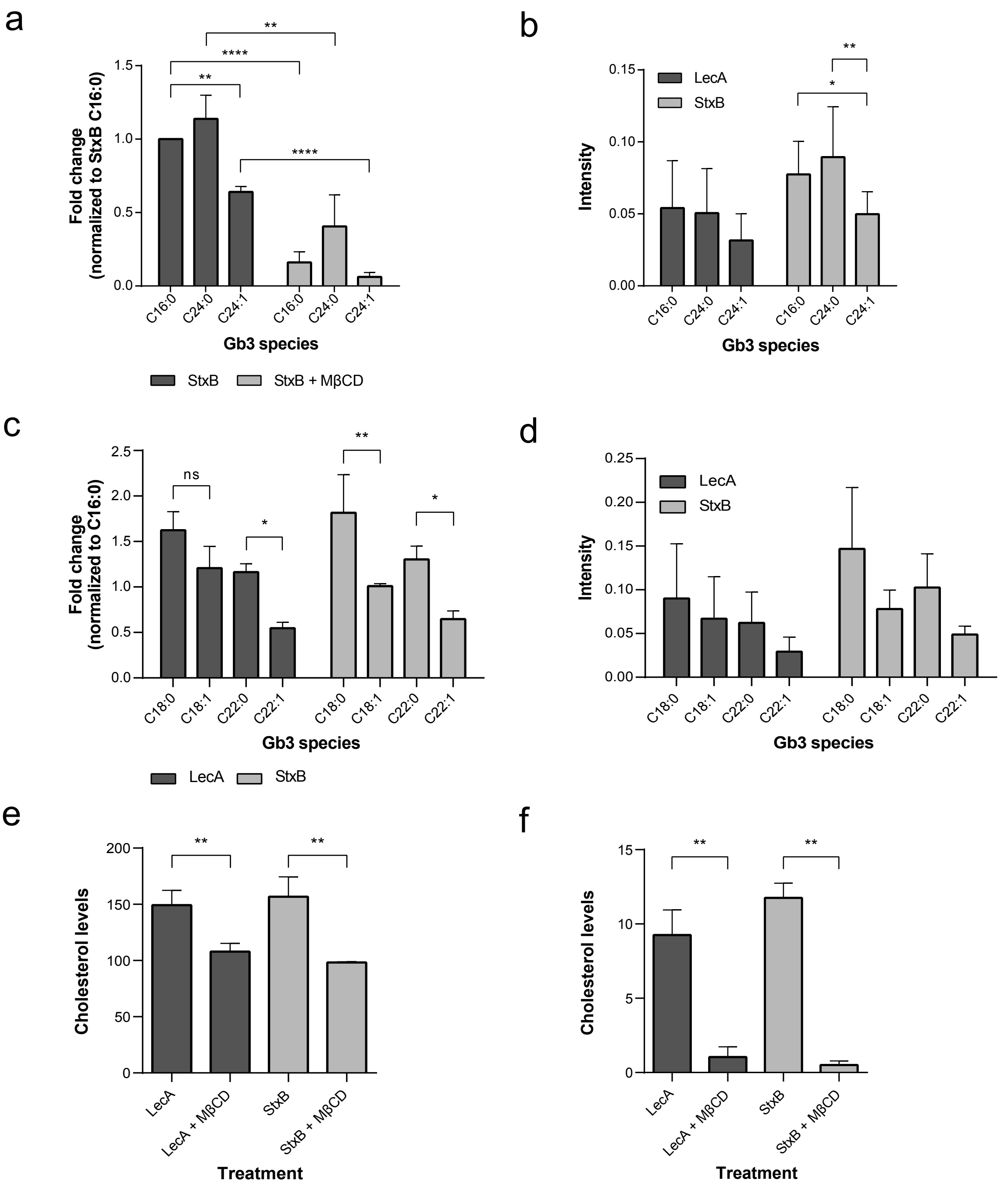


**Fig. S2** StxB demonstrates a predilection for saturated Gb3 species. **a** Pulled-down Gb3 species (normalized to input values and C16:0) of StxB-biotin demonstrated a preference for saturated over unsaturated Gb3 species. Significantly less Gb3 C16:0 and C24:1 was pulled-down in MβCD-treated cells stimulated with StxB-biotin. The Gb3 species C24:0 seemed less affected by MβCD treatment. Still, the unsaturated species C24:1 was pulled-down least. **b** Pulled-down Gb3 species of LecA- and StxB-biotin normalized to input levels but not to Gb3 C16:0 (in addition and contrast to Figs. 1B and S2A). Both lectins preferred saturated over unsaturated Gb3 species. **c,d** The pulled-down Gb3 species C18:0, C18:1, C22:0 and C22:1 of LecA- and StxB-biotin. Values are **c** normalized or **d** not normalized to C16:0. Also here, a preference for saturated species was visible. Remember, these Gb3 species are present in very low amounts within the plasma membrane of H1299 cells (Fig. 1a). **e**

Analysis of input lysates displayed a significant depletion of cholesterol upon MβCD treatment (10 mM). **f** The pulled-down cholesterol level was strongly reduced in cholesterol-depleted cells. For all panels: Bars display mean values of three biological replicates, error bars represent SD, *p<0.05, **p<0.01, p<****<0.0001 (For **a**, **b**, **c**, **d**: two-way ANOVA and Tukey's multiple comparisons tests; **e**, **f**: two-tailed unpaired t test).

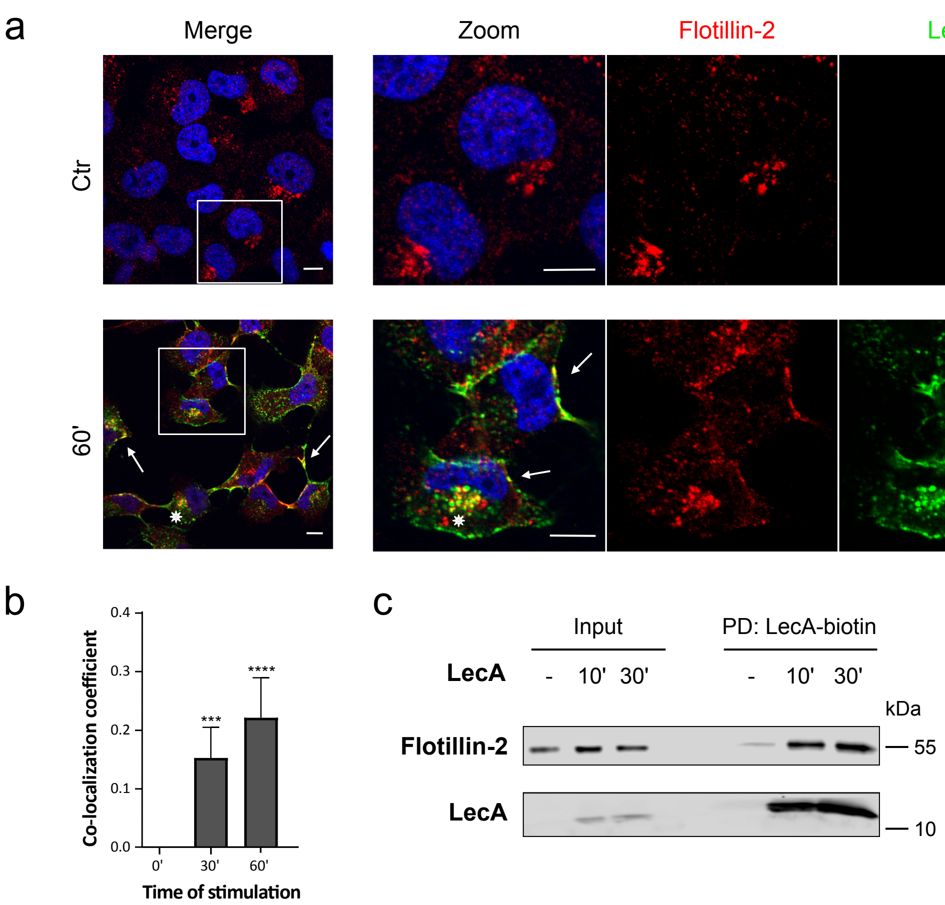


**Fig. S3** Flotillins are recruited to the plasma membrane upon LecA stimulation. **a** Fluorescent co-localization studies of flotillin-2 (red) and LecA (green) after 60 min of lectin stimulation. Nuclei were counterstained by DAPI. Framed areas were magnified. White arrows point at co-localization events at the plasma membrane, asterisks at perinuclear co-localization. Scale bar: 10 µm. **b** Co-localization between LecA and flotillin-2 was quantified by Mander's co-localization coefficient. Bars display mean values of three biological replicates, error bars represent SD, ***p<0.001, p<****<0.0001 (one-way ANOVA and Dunnett's multiple comparisons test). **c** Pull-down of LecA-biotin resulted in a time-dependent enrichment of flotillin-2.

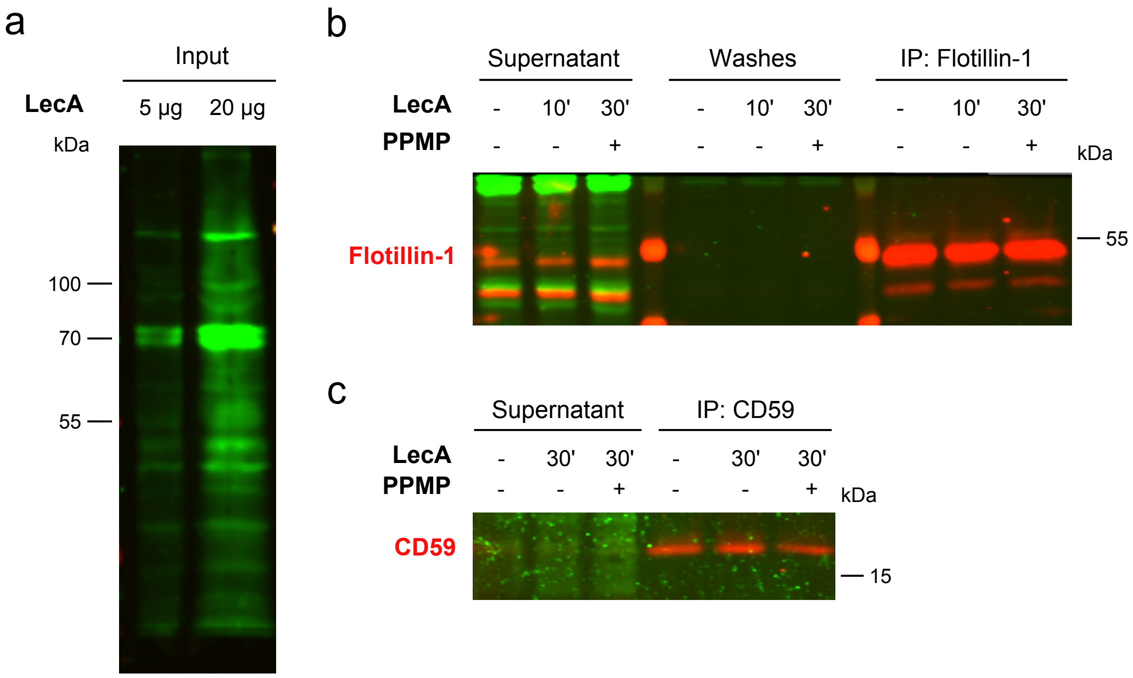


**Fig. S4** LecA does not directly bind to flotillin-1 or CD59. **a** WT H1299 lysates were subjected to SDS-PAGE and immunoblotting. The membrane was subsequently incubated with LecA-biotin for 60 min, followed by a 30 min-treatment of streptavidin-800 (green). Lectin blot shows proteins directly bound by LecA-biotin (via streptavidin). **b** Flotillin-1 co-immunoprecipitation of LecA-stimulated WT and PPMP-treated H1299 cells was conducted. Subsequently, lectin blot analysis of the obtained lysates was performed. No direct binding of LecA (via streptavidin, green) to flotillin-1 (red) in the eluates was detectable. **c** CD59 co-immunoprecipitation of LecA-stimulated WT and PPMP-treated H1299 cells was performed and analyzed in a lectin blot. No direct binding of LecA (via streptavidin, green) to CD59 (red) in the eluates was observed.

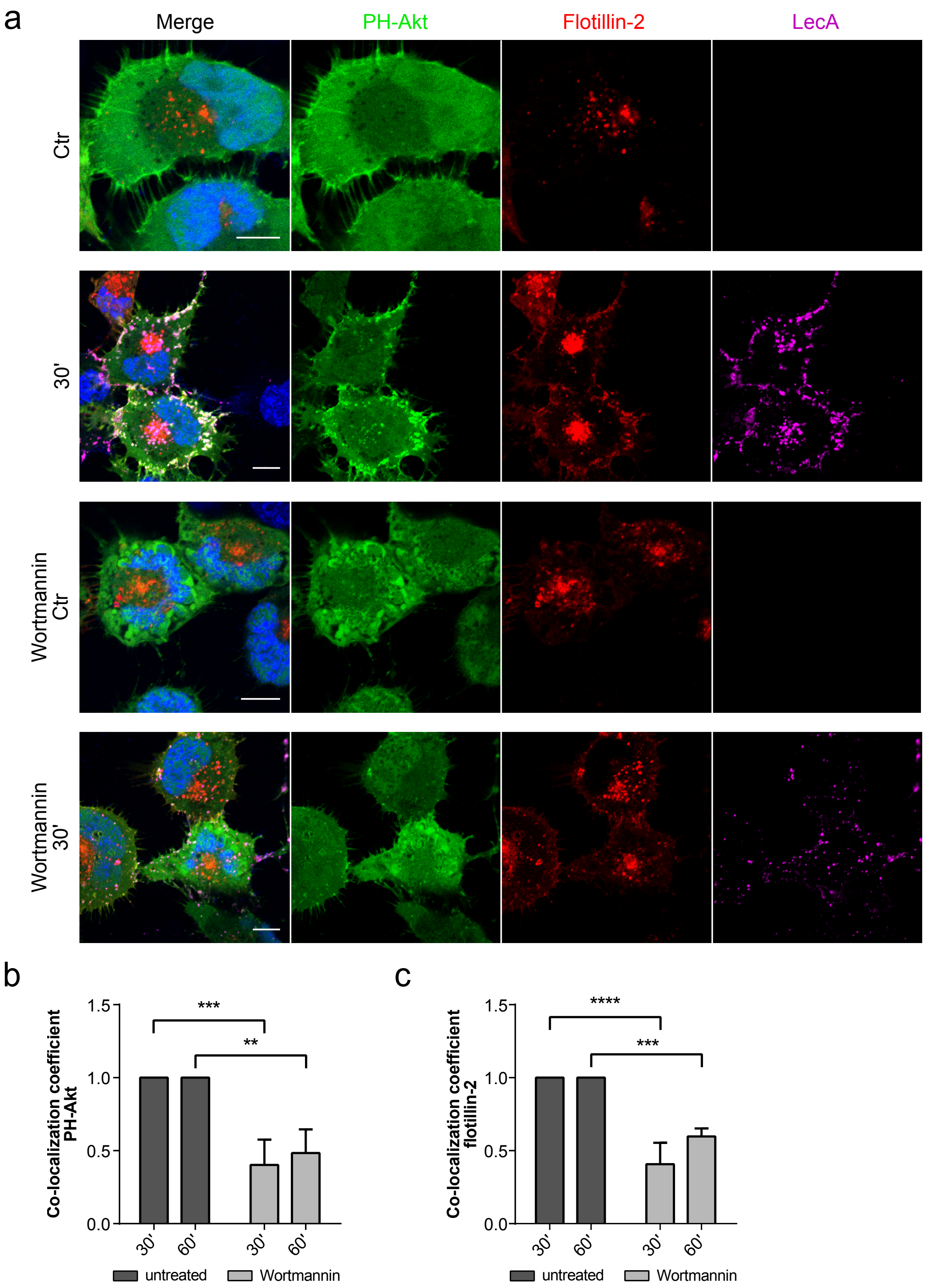


**Fig. S5** PI3-kinase inhibition partially prevents $PIP_3$ clustering and recruitment of flotillins upon LecA stimulation. **a** Confocal microscopy images of PH-Akt-GFP and flotillin-2-mCherry expressing H1299 cells exposed to fluorescent LecA. Lower two panels: Cells were pre-treated with 100 nM Wortmannin to inhibit PI3-kinase activity. Scale bar: 10 µm. **b** Co-localization of PH-Akt-GFP and LecA is depicted as Mander's co-localization coefficient normalized to the untreated conditions. **c** Mander's co-localization coefficient quantified between the fluorescent signals of flotillin-2-mCherry and LecA in comparison to the untreated conditions. Bars display mean values of three biological replicates, error bars represent SD, **p<0.01, ***p<0.001, ****p<0.0001 (two-way ANOVA and Tukey's multiple comparisons tests).

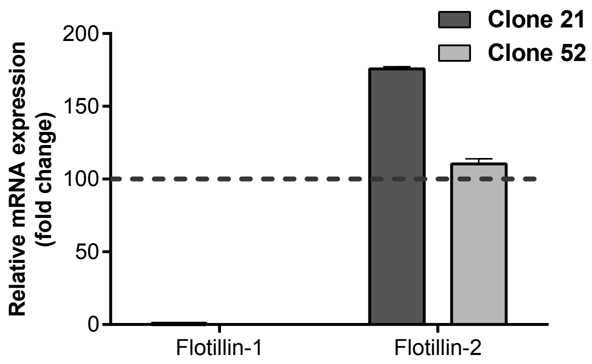


**Fig. S6** CRISPR-Cas9 knockout of flotillin-1. Flotillin-1 knockout efficiency of two clones was analyzed by qPCR and comparison of mRNA expression levels. For further experiments, clone 52 was chosen, as flotillin-2 mRNA expression appeared similar to WT levels. Bars display mean values of two biological replicates, error bars represent SD.

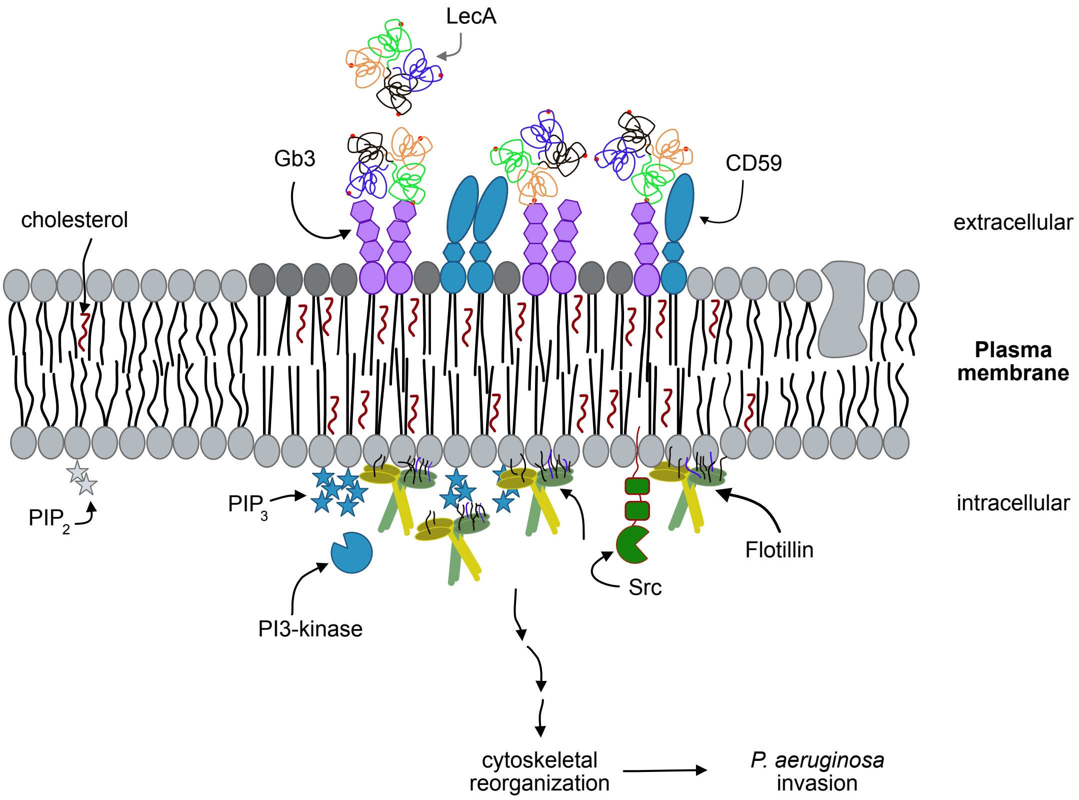


**Fig. S7** Potential model of the interactions of LecA with the host cell plasma membrane. LecA binds to its receptor, the GSL Gb3, inducing clustering of saturated Gb3 species and recruitment of the GPI-anchored protein CD59 within the extracellular leaflet. At the intracellular leaflet PI3-kinases phosphorylate $PIP_2$ to $PIP_3$, which influences the recruitment of flotillins to the LecA-induced plasma membrane domain. The signal may be transduced from the extracellular to the intracellular site by transbilayer coupling between long fatty acyl chains of Gb3 and CD59 on the one hand and $PIP_3$ on the other hand. Small GTPases like Rac1 and Src family kinases are known to mediate cytoskeletal reorganization. The formation of the LecA plasma membrane domain might prime the host cell for an efficient uptake of *P. aeruginosa*. Components are not drawn to scale.

**Table S1** Sequences of used siRNAs, primers for qPCR and sgRNAs for CRISPR-Cas knockout experiments.

| Name | Sequence | Company |
|---|---|---|
| Flotillin-2 siRNA target sequences | UUGAGGUUGUGCAGCGCAA, AGGCAGAGGCUGAGCGGAU, GACAAGGAGCUCAUCGCUA, CGGAAGCGGCAGUCAUCGA | Dharmacon, Horizon |
| CD59 siRNA target sequences | AAAAUGAGCUAACGUACUA, GCAAGAAGGACCUGUGUAA, CAGAAUAGCUUGGUCCUUA, GUGUGAGGCUGGCGCGAUU | Dharmacon, Horizon |
| Flotillin-1 primer for qPCR | TCCAGCCTGAACCATGTTTT ATTGAGGGTCAGTGTGTTGA | Sigma-Aldrich |
| Flotillin-2 primer for qPCR | GGACGTGTATGACAAAGTGG GTGTCTGCCATGAACTTCAC | Sigma-Aldrich |
| GAPDH primer for qPCR | CAAAATCAAGTGGGGCGATG TTGCTGATGATCTTGAGGCT | Sigma-Aldrich |
| Flotillin-1 sgRNA1 for CRISPR-Cas | GGGGTGCTTAGTCCGTTGCG Pam sequence: GGG | Eurofins |
| Flotillin-1 sgRNA2 for CRISPR-Cas | GAAGGCTGACATGGGACCGC Pam sequence: TGG | Eurofins |

**Table S2** The MS protein hits: The MS protein analysis revealed a list of hits mainly including cytoskeleton and cytoskeletal-related components as well as small GTPases. Hits are ordered according to abundance. Listed are only proteins detected in both replicates and present with an emPAI >0.01%.

| **Name** | **Accession** | **LecA 5'** | **LecA 15'** |
|---|---|---|---|
| Vimentin | VIME_HUMAN | + | + |
| Actin, cytoplasmic 1 | ACTB_HUMAN | - | + |
| Actin, alpha cardiac muscle 1 | ACTC_HUMAN | - | + |
| Myosin-9 | MYH9_HUMAN | - | + |
| Ubiquitin-40S ribosomal protein S27a | RS27A_HUMAN | - | + |
| Tubulin beta-6 chain | TBB6_HUMAN | - | + |
| CD59 glycoprotein | CD59_HUMAN | + | + |
| Ras-related C3 botulinum toxin substrate 1 | RAC1_HUMAN | + | + |
| Rac GTPase-activating protein 1 | RGAP1_HUMAN | + | + |
| Tyrosine-protein kinase Yes | YES_HUMAN | - | + |
| F-actin-capping protein subunit alpha-1 | CAZA1_HUMAN | - | + |
| RCC1 and BTB domain-containing protein 1 | RCBT1_HUMAN | - | + |
| RCC1 and BTB domain-containing protein 2 | RCBT2_HUMAN | - | + |
| F-actin-capping protein subunit beta | CAPZB_HUMAN | - | + |
| Prohibitin-2 | PHB2_HUMAN | - | + |
| DDB1- and CUL4-associated factor 13 | DCA13_HUMAN | - | + |
| Flotillin-1 | Flot1_HUMAN | - | + |
| Sequestosome-1 | SQSTM_HUMAN | - | + |
| NKAP-like protein | NKAPL_HUMAN | + | + |
| GPI transamidase component PIG-T | PIGT_HUMAN | - | + |
| Tyrosine-protein kinase BAZ1B | BAZ1B_HUMAN | - | + |
| Scaffold attachment factor B1 | SAFB1_HUMAN | - | + |
| Supervillin | SVIL_HUMAN | - | + |
